# Analytical solution for a hybrid Logistic-Monod cell growth model in batch and CSTR culture

**DOI:** 10.1101/847939

**Authors:** Peng Xu

## Abstract

Monod and Logistic growth models have been widely used as basic equations to describe cell growth in bioprocess engineering. In the case for Monod equation, specific growth rate is governed by a limiting nutrient, with the mathematical form similar to the Michaelis-Menten equation. In the case for Logistic equation, specific growth rate is determined by the carrying capacity of the system, which could be growth-inhibiting factors (i.e. toxic chemical accumulation) other than the nutrient level. Both equations have been found valuable to guide us build structure-based kinetic models to analyze fermentation process and understand cell physiology. In this work, we present a hybrid Logistic-Monod growth model, which accounts for multiple growth-dependence factors including both the limiting nutrient and the carrying capacity of the system. Coupled with substrate consumption and yield coefficient, we present the analytical solutions for this hybrid Logistic-Monod model in both batch and CSTR culture. Under high biomass yield (*Y*x/s) conditions, the analytical solution for this hybrid model is approaching to the Logistic equation; under low biomass yield condition, the analytical solution for this hybrid model converges to the Monod equation. This hybrid Logistic-Monod equation represents the cell growth transition from substrate-limiting condition to growth-inhibiting condition, which could be adopted to accurately describe the multi-phases of cell growth and may facilitate kinetic model construction, bioprocess optimization and scale-up in industrial biotechnology.

---

In 1949, French microbiologist Dr. Jacques Monod (who was also a Nobel Laureate in 1965, best known for his discovery of Lac operon), provided a quantitative description between bacterial growth rate and the concentration of a limiting substrate (glucose) (Monod, 1949). This equation (**Eqn. 1**) takes the mathematical form of Michaelis-Menten equation, where the substrate saturation constant (*K*s) and the maximal specific cell growth rate (*μ*_max_) could be graphically determined by the Lineweaver–Burk double-reciprocal plot (Lineweaver & Burk, 1934). Under limiting nutrient conditions, (*S* << *K*s), cell growth follows first order kinetics and the specific growth rate is proportional to the nutrient level. Under unlimited nutrient conditions (*S* → +∞), cells could reach their maximal growth potential (the cell is therefore saturated by the substrate) and follow zeroth order kinetics. The specific growth rate follows a monotonically increasing pattern as we increase the concentration of the limiting nutrient (*S*).

The Logistic equation (**Eqn. 2**), was first introduced by the UK sociologist Thomas Malthus to describe the “the law of population growth” at the end of 18^th^ century. This model later was formulated and derived by the Belgian mathematician Pierre François Verhulst to describe the self-limiting growth of a biological population in 1838. With little self-limiting factor (*X* → 0), the population attains the maximal grow rate (*μ*_max_). As the cell growth, the population starts inhibiting themselves (could be considered as a negative auto-regulation loop). With sufficient self-limiting factors (*X* → *X*m), the population reaches the carrying capacity of the system and the growth rate approaches to zero. The specific growth rate follows a linearly decreasing pattern as the cell population (*X*) increases.

Both Monod and Logistic model have been used extensively to analyze fermentation process and study microbial consortia interactions. For example, an expanded form of Monod equation was proposed to account for product, cell, and substrate inhibitions (Han & Levenspiel, 1988; Levenspiel, 1980; Luong, 1987). When the Monod equation was coupled with the Luedeking-Piret equation (Robert Luedeking, 1959), analytical solutions for cell growth, substrate consumption and product formation could be derived (Garnier & Gaillet, 2015). A square-root boundary between cell growth rate and biomass yield has been proposed (Wong, Tran, & Liao, 2009). Coupled Monod equations were applied to describe the complicated predator-prey (oscillatory) relationship between *Dictyostelium discoideum* and *Escherichia coli* in Chemostat (Tsuchiya, Drake, Jost, & Fredrickson, 1972). Much earlier than the Monod equation, the Logistic growth was used by the American biophysicist Alfred J. Lotka and the Italian mathematician Vito Volterra to describe the famous Lotka–Volterra predator-prey ecological model (Lotka, 1926; Volterra, 1926). More interestingly, the solutions of the discrete Logistic growth model were elegantly analyzed by the Australian ecologist Robert May (Baron May of Oxford) in early 1970s. It was discovered that complex dynamic behaviors could arise from this simple Logistic equation, ranging from stable points, to bifurcating stable cycles, to chaotic fluctuations, all depending on the initial parameter conditions (May, 1976). Both models benefit us to analyze the microbial process and explore unknown biological phenomena.

To account for both substrate-limiting and self-inhibiting factors, herein we propose a hybrid Logistic-Monod model (**Eqn. 3**) by simply multiplying the Monod equation with the self-inhibiting factor (1 − *X*/*Xm*). We believe this model could better describe the transition of cell growth from substrate-limiting phases to self-inhibiting phases. Analytical solutions for this hybrid model will be derived for both batch and CSTR cultivation mode in the following section.

We will first look the analytical solution for the Monod equation. By coupling Monod equation (**Eqn. 1**) with substrate consumption rate (**Eqn. 4**) and yield coefficient (*Y*xs), the implicit form of the analytical solution for cell growth (*X*, **Eqn. 5**) and substrate (*S*, **Eqn. 6**) could be easily solved by separation of variables or Laplace transformation. A typical Monod-type kinetics was plotted for batch culture (**Fig. 1a**). It should be noted that the initial conditions are prescribed as *S* = *S*_0_ and *X* = *X*_0_ at the beginning of cultivation (*t* = 0).

**Fig. 1.**
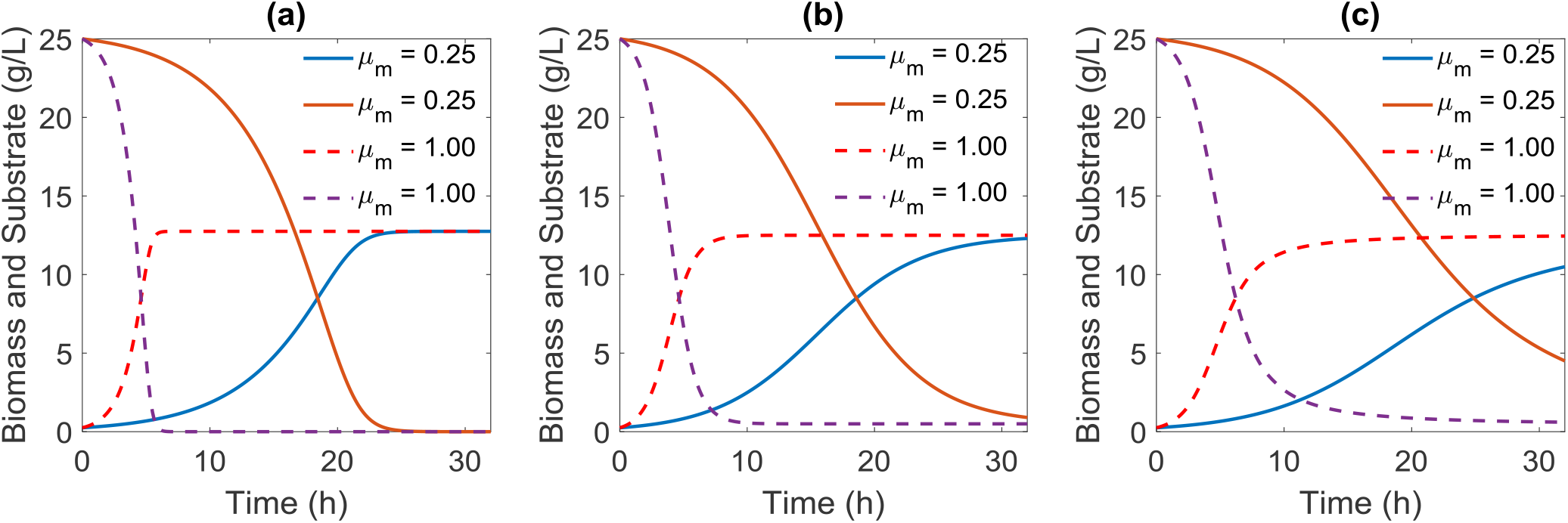
Analytical solutions for three growth models in batch culture. (**a**) Monod growth model; (**b**) Logistic growth model; and (**c**) the hybrid Logistic-Monod growth model. All the three models have the same parameter settings (*X*_m_=12.5 g/L; *Y*_xs_=0.5 g/g; *K*_S_=4 g/L; *S*_0_ = 25 g/L; and *X*_0_ = 0.25 g/L) except for *μ*_m_. The dotted line represents high growth potential (*μ*_m_); and the solid line represents low growth potential (*μ*_m_).

In the case for Logistic model, we could also arrive the analytical solutions for cell growth (*X*, **Eqn. 7**) and substrate (*S*, **Eqn. 8**) by separation of variables or Laplace transformation, when the Logistic equation (**Eqn. 2**) is coupled with substrate consumption kinetics (**Eqn. 4**). It should be noted that the cell growth is independent of substrate consumption in the Logistic model, but the substrate will deplete proportionally with cell growth (**Fig. 1b**). Due to the simplicity of the Logistic equation, we could arrive the explicit solution for cell growth (*X*) and substrate (*S*).

Similarly, by coupling **Eqn.3** with **Eqn. 4**, the implicit solutions for the hybrid Logistic-Monod equations (**Eqn. 3**) could be derived analytically with the aid of the symbolic computation package of MATLAB. This hybrid Logistic-Monod model (**Eqn. 3**) retains the elementary differential equation norm and should be solved analytically by either separation of variables or Laplace transformation, despite the derivation process will be trivial. The exact solution for cell growth (*X*, **Eqn. 9**) and substrate (*S*, **Eqn. 10**) takes a much-complicated form, due to the fact that this hybrid model accounts for both nutrient-limiting factors (*K*s) and self-inhibiting factors (*X*_m_). Cell growth and substrate consumption take the sigmoidal pattern (**Fig. 1c**), and the biomass formation and substrate depletion are approaching to the Monod model in the short-time regime (which is a nutrient-limiting phase), but shift toward the Logistic model in the long-time regime (which is self-inhibiting phase).

We next will explore the steady state solutions of three growth models in CSTR culture. Based on mass balance and the substrate concentration in the feeding stream (*S*_F_), we could list the mass balance for cell growth (**Eqn. 11**) and substrate consumption (**Eqn. 12**). When the CSTR mass balance equations (**Eqn. 11** and **Eqn. 12**) are coupled with the Monod growth kinetics **(Eqn. 1**), it is easy to arrive the steady state substrate and cell concentration in the CSTR (**Eqn. 13** and **Eqn. 14**), which has been widely taught in Biochemical engineering or Bioprocess engineering textbooks. As the dilution rate increases, the substrate concentration increases with decreasing cell concentration at the outlet flow of the CSTR (**Fig. 2a**). Similarly, when the mass balance equations (**Eqn. 11** and **Eqn. 12**) are coupled with the Logistic growth kinetics (**Eqn. 2**), the steady state solutions for substrate and biomass could be derived analytically (**Eqn. 15** and **Eqn. 16**). As the dilution rate increases, the substrate concentration linearly increases accompanying with proportionally decreased cell concentration at the outlet flow of the CSTR (**Fig. 2b**). Finally, for the hybrid Logistic-Monod model, we could also derive the steady state solutions for the substrate and biomass concentration (**Eqn. 17** and **Eqn. 18**, **Fig. 2c**), when the CSTR mass balance equations (**Eqn. 11** and **Eqn. 12**) are coupled with the hybrid Logistic-Monod model (**Eqn. 3**). Plotting all three models together (**Fig. 2**), it is clear that the solution for the hybrid Logistic-Monod model (**Fig. 2c**) is approaching to the Monod model (**Fig. 2a**) under low-biomass-yield (*Y*_xs_= 0.3 g/g) regime, indicating a nutrient-limiting condition; and the steady state solution (**Fig. 2c**) will shift toward the Logistic model (**Fig. 2b**) under high-biomass-yield (*Y*_xs_ = 0.8 g/g) regime, indicating a self-inhibiting condition. In practical, the maximal cell density *X*_m_ should take a value below *S*_0_*Y*_xs_ (*X*_m_ ≤ *S*_0_*Y*_xs_ or *X*_m_ ≤ *S*_F_*Y*_xs_) for both the batch and CSTR culture, assuming that all the substrate could be converted to biomass.

**Fig. 2.**
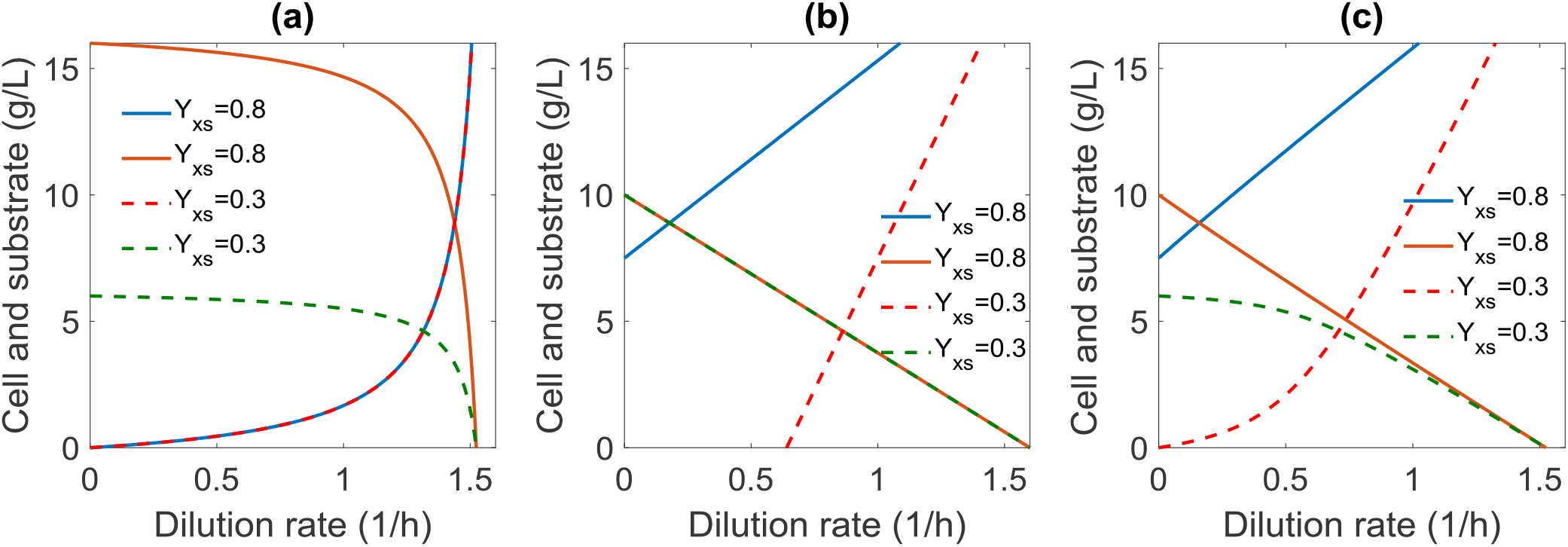
Analytical solutions for three growth models in CSTR culture. (**a**) Monod growth model; (**b**) Logistic growth model; and (**c**) the hybrid Logistic-Monod growth model. All the three models have the same parameter settings (*μ*_m_ = 1.6 hr^−1^; *X*_m_=10 g/L; *K*_S_=1 g/L; and *S*_F_ = 20 g/L) except for *Y*_XS_. The dotted line represents the low-biomass-yield (*Y*_xs_) regime, and solid line represents high-biomass-yield regime. In Fig. 2c, the hybrid Logistic-Monod model captures the feature for both nutrient-limited conditions (the dotted lines represent a typical Monod growth as shown in Fig. 2a) and the self-inhibitory conditions (the solid lines represent a typical Logistic growth as shown in Fig. 2b).

Taken together, this hybrid Logistic-Monod model represents cellular physiology transition from nutrient-limiting conditions to self-inhibiting conditions. It expands the classical Monod and Logistic growth model, and should find broad applications to analyze fermentation process (Xu, Qiao, & Stephanopoulos, 2017) and study microbial consortia interactions. With the proposed Logistic-Monod equation, the product formation kinetics could also be solved, when the above models are coupled with Gaden equation (Gaden Jr, 1959). By introducing the carrying capacity concept into the classical Monod equation, this hybrid Logistic-Monod Model should also find applications in gene circuits engineering to study growth-dependent gene expression pattern (Aris et al., 2019) or burden effects that are related with gene overexpression (Ceroni, Algar, Stan, & Ellis, 2015; Ceroni et al., 2018). More importantly, we present the analytical solutions for this hybrid Logistic-Monod growth model in both batch and CSTR culture. These analytical solutions may be integrated with process control software (Shirsat et al., 2015; Zhang, Del Rio-Chanona, Petsagkourakis, & Wagner, 2019) to improve our prediction accuracy for both the bench-top or pilot-scale bioreactors, and facilitate us to design more efficient microbial process for various industrial applications.

## Computational Methods

Matlab R2017b was used as the computational platform on a Windows 7 professional operation system. The CPU processor is Intel Core i3-6100 with 3.70 GHz. The installed memory (RAM) is 4.0 GHz. Matlab symbolic language package coupled with LaTex makeup language were used to derive and output the solution. Matlab implicit function fplot was used to draw most of the solutions for Fig. 1 and Fig. 2. Matlab code has been compiled into a supplementary file, has been uploaded to the journal website.

**Table 1.**
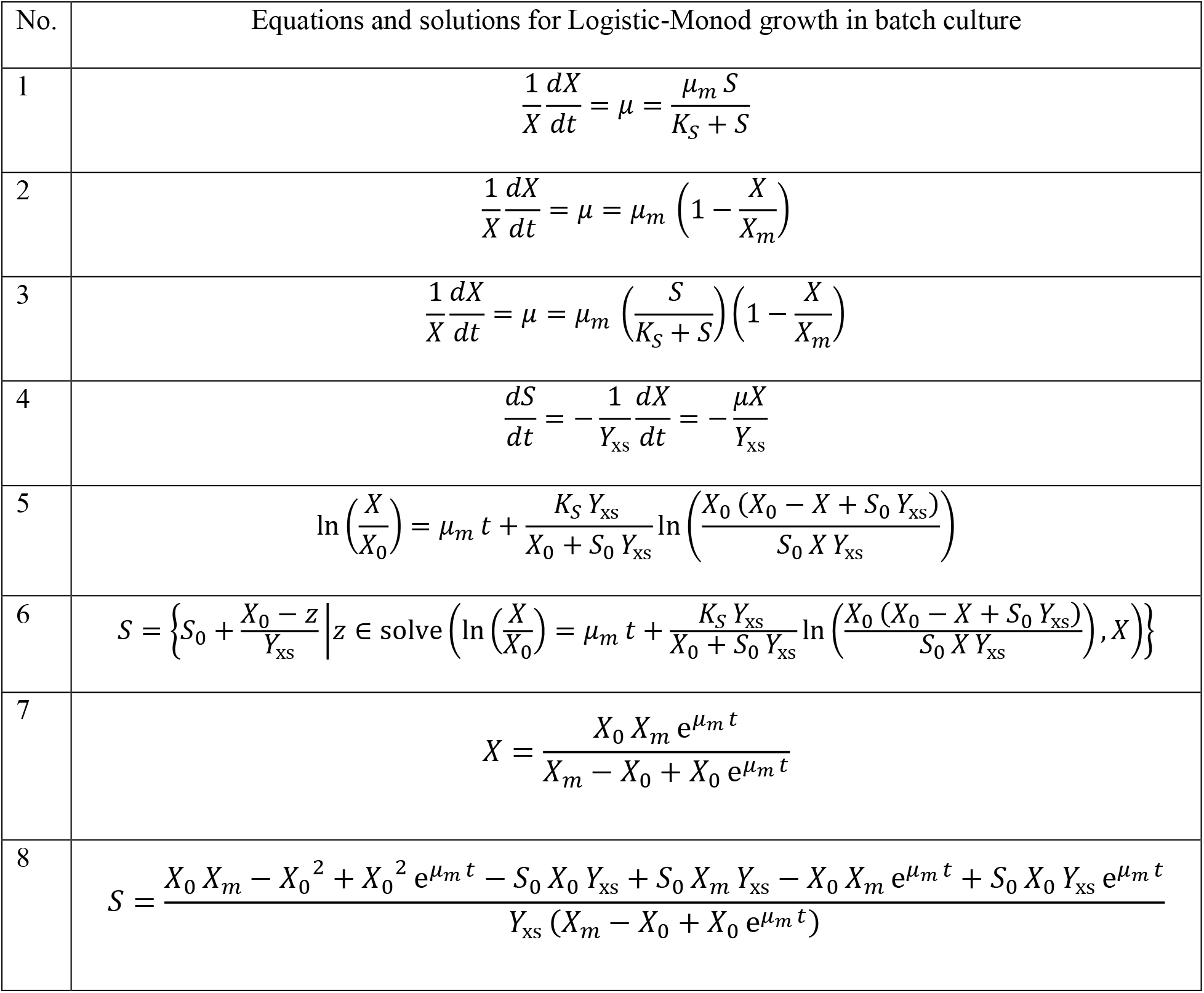
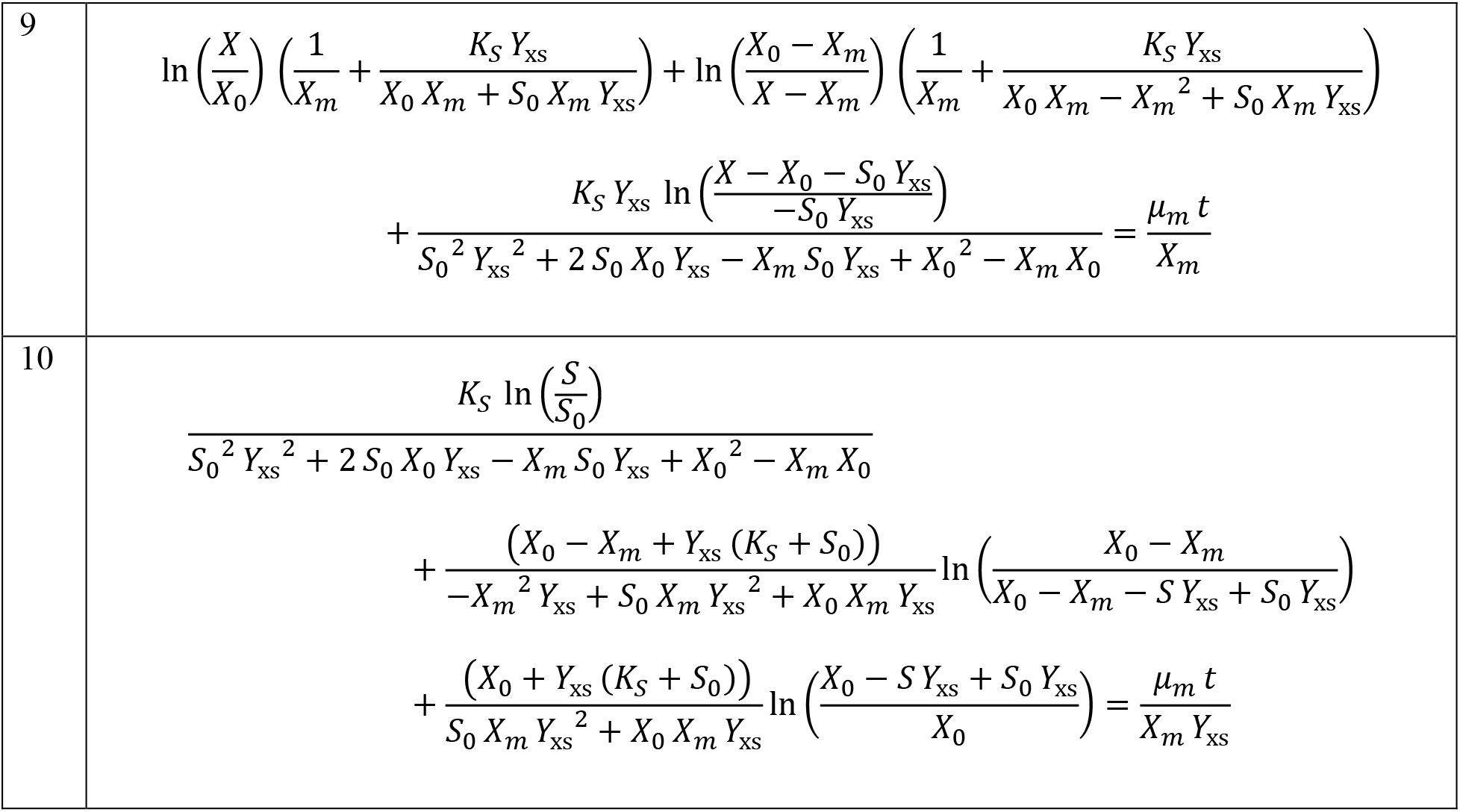
List of equations and solutions for Logistic-Monod growth in batch culture

**Table 2.**
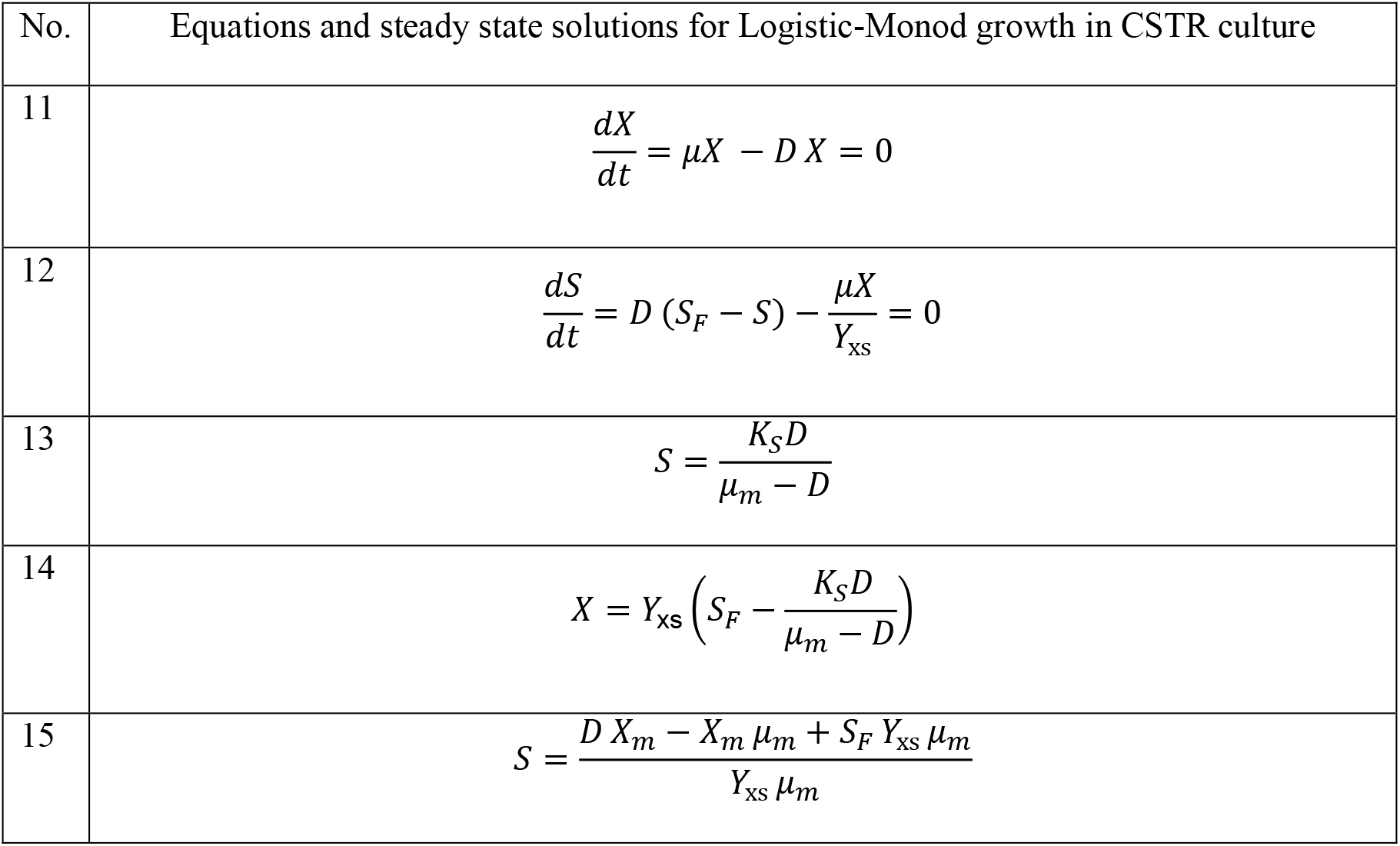
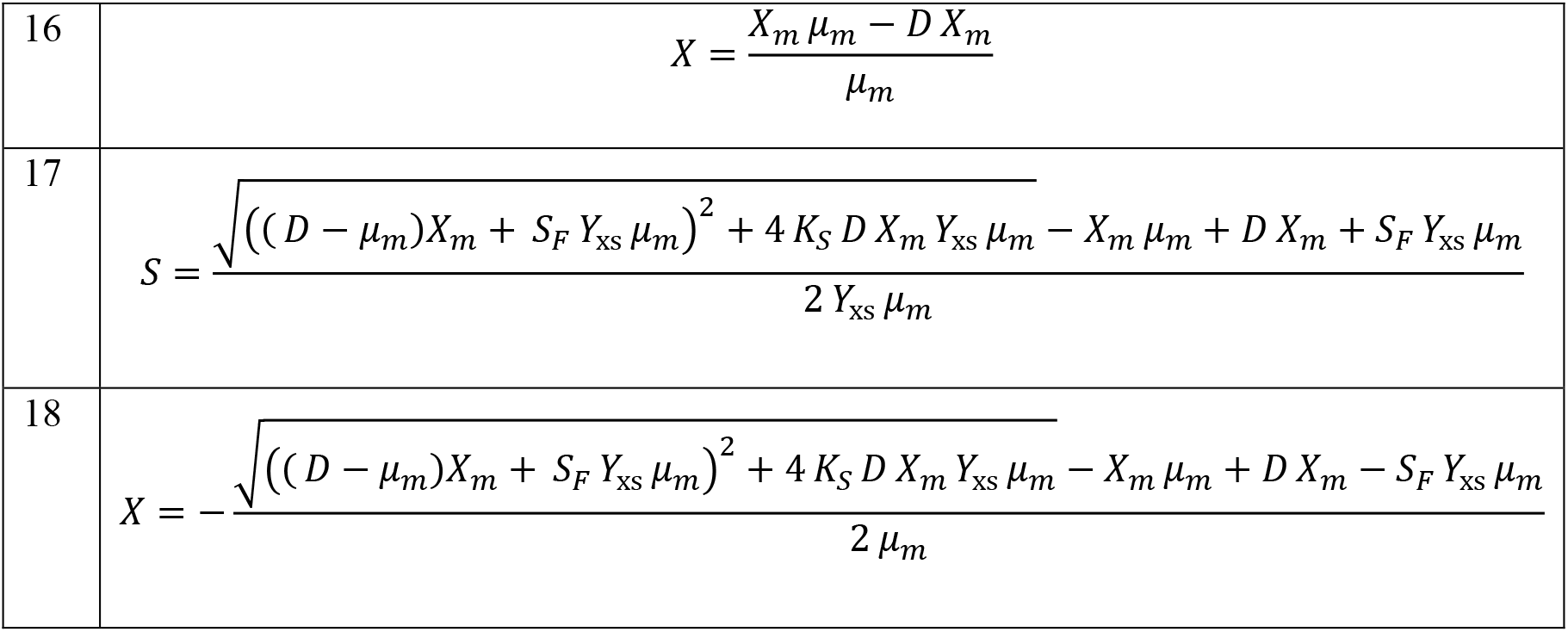
List of equations and solutions for Logistic-Monod growth in CSTR culture

## Supporting information

Supplemental Matlab code

## Acknowledgments

Dr. Xu would like to acknowledge the Bill and Melinda Gates Foundation (OPP1188443) and National Science Foundation under grant number 1805139 for financially supporting this project. Dr. Xu would also like to thank the previous mentors that may allow himself to dive into the classical biochemical engineering arena.

---

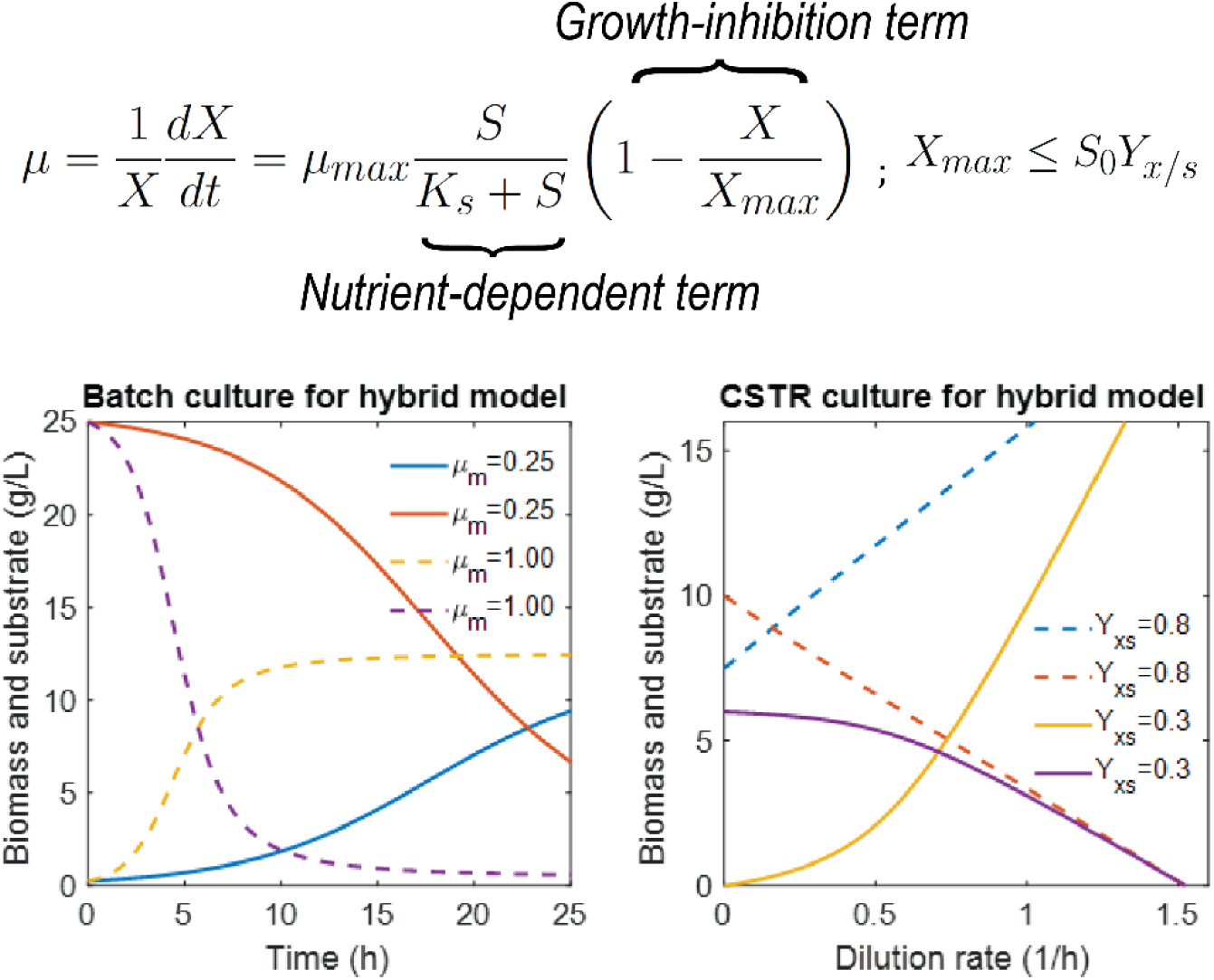
Graphic Abstract

