## Supplemental Matlab code for "Analytical solution for a hybrid Logistic-Monod cell growth model in batch and CSTR culture"

### Running title: Analytical solution for hybrid Logistic-Monod model

Peng Xu*

Department of Chemical, Biochemical and Environmental Engineering, University of Maryland

Baltimore County, Baltimore, MD 21250

 (PX)

### 1. Matlab code to solve the Monod growth model in a batch fermentation

```
>> syms mu_m K_S X(t) X0 S0 Y_xs S(t)

eqn5 = diff(X,t) == mu_m*(S0-(X-X0)/Y_xs)*X/(K_S+ S0-(X-X0)/Y_xs)

cond1 = X(0) == X0;

solX =dsolve(eqn5,cond1);

eqn5(t) =


diff(X(t), t) == (mu_m*X(t)*(S0 + (X0 - X(t))/Y_xs))/(K_S + S0 + (X0 - X(t))/Y_xs)

>> latex(eqn5)


ans =
```

$$\frac{\partial}{\partial t}X(t) = \frac{\mu_m\,X(t)\,\left(S_0 + \frac{X_0 - X(t)}{Y_{\mathrm{xs}}}\right)}{K_S + S_0 + \frac{X_0 - X(t)}{Y_{\mathrm{xs}}}}$$

```
solX =


solve(log(X) - log(X0) - mu_m*t - (2*K_S*Y_xs*atanh((X0 - 2*X + S0*Y_xs)/(X0 + S0*Y_xs)))/(X0 + S0*Y_xs) - (2*K_S*Y_xs*atanh((X0 - S0*Y_xs)/(X0 + S0*Y_xs)))/(X0 + S0*Y_xs), X)


>> latex(solX)

ans =
```

$$\mathrm{solve}\left(\ln(X) - \frac{2\,K_S\,Y_{\mathrm{xs}}\,\mathrm{atanh}\left(\frac{X_0 - 2\,X + S_0\,Y_{\mathrm{xs}}}{X_0 + S_0\,Y_{\mathrm{xs}}}\right)}{X_0 + S_0\,Y_{\mathrm{xs}}} = \ln(X_0) + \mu_m\,t + \frac{2\,K_S\,Y_{\mathrm{xs}}\,\mathrm{atanh}\left(\frac{X_0 - S_0\,Y_{\mathrm{xs}}}{X_0 + S_0\,Y_{\mathrm{xs}}}\right)}{X_0 + S_0\,Y_{\mathrm{xs}}}, X\right)$$

```
>> syms mu_m K_S X X0 S0 Y_xs S

>> eqn1 = log(X/X0) ==  mu_m*t + K_S*Y_xs/(X0+S0*Y_xs)*log((X0/(S0*Y_xs))*(X0-X+S0*Y_xs)/X);

>> latex(eqn1)
```

$$\ln\left(\frac{X}{X_0}\right) = \mu_m\,t + \frac{K_S\,Y_{\mathrm{xs}}}{X_0 + S_0\,Y_{\mathrm{xs}}}\ln\left(\frac{X_0\,(X_0 - X + S_0\,Y_{\mathrm{xs}})}{S_0\,X\,Y_{\mathrm{xs}}}\right)$$

```
>> solS = S0-(solX-X0)/Y_xs
```

solS =

{S0 + (X0 - z)/Y_xs | z in solve(log(X) - (2*K_S*Y_xs*atanh((- 2*X + X0 + S0*Y_xs)/(X0 + S0*Y_xs)))/(X0 + S0*Y_xs) == log(X0) + mu_m*t + (2*K_S*Y_xs*atanh((X0 - S0*Y_xs)/(X0 + S0*Y_xs)))/(X0 + S0*Y_xs), X)}

>> latex(solS)

ans =

$$\left\{S_0+\frac{X_0-z}{Y_{\mathrm{xs}}}\;\middle|\; z\in \mathrm{solve}\left(\ln(X)-\frac{2\,K_S\,Y_{\mathrm{xs}}\,\mathrm{atanh}\left(\frac{X_0-2\,X+S_0\,Y_{\mathrm{xs}}}{X_0+S_0\,Y_{\mathrm{xs}}}\right)}{X_0+S_0\,Y_{\mathrm{xs}}}=\ln(X_0)+\mu_m\,t+\frac{2\,K_S\,Y_{\mathrm{xs}}\,\mathrm{atanh}\left(\frac{X_0-S_0\,Y_{\mathrm{xs}}}{X_0+S_0\,Y_{\mathrm{xs}}}\right)}{X_0+S_0\,Y_{\mathrm{xs}}},X\right)\right\}$$

### 2. Matlab code to solve the Logistic growth model in a batch fermentation

```
>> syms mu_m X_m X(t) X0 S0 Y_xs S(t)
>> eqn1 = diff(X,t) == mu_m*X*(1-X/X_m)
>> eqn2 = diff(S,t) == - mu_m*X*(1-X/X_m)/Y_xs;
>> cond1 = X(0) == X0;
>> cond2 = S(0) == S0;
>> eqns = [eqn1, eqn2]
>> conds =[cond1, cond2]
>> solLogistic =dsolve(eqns,conds);
>> eqn3 = X(t) == simplify(solLogistic.X);
>> eqn4 = S(t) == simplify(solLogistic.S);
>> latex(eqn1)
```

ans =

$$\frac{\partial}{\partial t}X(t)=-\mu_m\,X(t)\,\left(\frac{X(t)}{X_m}-1\right)$$

>> latex(eqn2)

ans =

$$\frac{\partial}{\partial t} S(t) = \frac{\mu_m X(t) \left(\frac{X(t)}{X_m} - 1\right)}{Y_{xs}}$$

>> latex(eqn3)

ans =

$$X(t) = \frac{X_0 X_m e^{\mu_m t}}{X_m - X_0 + X_0 e^{\mu_m t}}$$

>> latex(eqn4)

ans =

$$S(t) = \frac{X_0 X_m - {X_0}^2 + {X_0}^2 e^{\mu_m t} - S_0 X_0 Y_{xs} + S_0 X_m Y_{xs} - X_0 X_m e^{\mu_m t} + S_0 X_0 Y_{xs} e^{\mu_m t}}{Y_{xs} (X_m - X_0 + X_0 e^{\mu_m t})}$$

### 3. Matlab code to solve the Logistic-Monod Model for Batch culture

```
>> syms mu_m K_S X_m X(t) X0 S0 Y_xs S(t)
>> eqn6 = diff(X,t) == mu_m*(S0-(X-X0)/Y_xs)*(1-X/X_m)*X/(K_S+ S0-(X-X0)/Y_xs)
>> cond1 = X(0) == X0;
>> solX =simplify(dsolve(eqn6,cond1));
>> latex(eqn6)
ans =
```

$$\frac{\partial}{\partial t} X(t) = -\frac{\mu_m X(t) \left(\frac{X(t)}{X_m} - 1\right) \left(S_0 + \frac{X_0 - X(t)}{Y_{xs}}\right)}{K_S + S_0 + \frac{X_0 - X(t)}{Y_{xs}}}$$

```
>> solX

solX =

log(X/X0)*(1/X_m + (K_S*Y_xs)/(X0*X_m + S0*X_m*Y_xs)) + log((X0 - X_m)/(X -
X_m))*(1/X_m + (K_S*Y_xs)/(X0*X_m - X_m^2 + S0*X_m*Y_xs)) + (K_S*Y_xs*log((X - X0 -
```

S0*Y_xs)/(-S0*Y_xs)))/(S0^2*Y_xs^2 + 2*S0*X0*Y_xs - X_m*S0*Y_xs + X0^2 - X_m*X0) - (mu_m*t)/X_m

>> latex(solX)

ans =

$$\ln\left(\frac{X}{X_0}\right)\,\left(\frac{1}{X_m}+\frac{K_S\,Y_{\mathrm{xs}}}{X_0\,X_m+S_0\,X_m\,Y_{\mathrm{xs}}}\right)+\ln\left(\frac{X_0-X_m}{X-X_m}\right)\,\left(\frac{1}{X_m}+\frac{K_S\,Y_{\mathrm{xs}}}{X_0\,X_m-{X_m}^2+S_0\,X_m\,Y_{\mathrm{xs}}}\right)+\frac{K_S\,Y_{\mathrm{xs}}\,\ln\left(\frac{X-X_0-S_0\,Y_{\mathrm{xs}}}{-S_0\,Y_{\mathrm{xs}}}\right)}{{S_0}^2\,{Y_{\mathrm{xs}}}^2+2\,S_0\,X_0\,Y_{\mathrm{xs}}-X_m\,S_0\,Y_{\mathrm{xs}}+{X_0}^2-X_m\,X_0}=\frac{\mu_m\,t}{X_m}$$

$$\mathrm{solve}\left(\frac{\ln(X)\,\left(X_0+K_S\,Y_{\mathrm{xs}}+S_0\,Y_{\mathrm{xs}}\right)}{X_m\,\left(X_0+S_0\,Y_{\mathrm{xs}}\right)}-\frac{\ln(X-X_m)\,\left(X_0-X_m+K_S\,Y_{\mathrm{xs}}+S_0\,Y_{\mathrm{xs}}\right)}{X_m\,\left(X_0-X_m+S_0\,Y_{\mathrm{xs}}\right)}+\frac{K_S\,Y_{\mathrm{xs}}\,\ln(X-X_0-S_0\,Y_{\mathrm{xs}})}{\left(X_0+S_0\,Y_{\mathrm{xs}}\right)\,\left(X_0-X_m+S_0\,Y_{\mathrm{xs}}\right)}=\frac{\mu_m\,t}{X_m}-\frac{\ln(X_0-X_m)\,\left(X_0-X_m+K_S\,Y_{\mathrm{xs}}+S_0\,Y_{\mathrm{xs}}\right)}{X_m\,\left(X_0-X_m+S_0\,Y_{\mathrm{xs}}\right)}+\frac{\ln(X_0)\,\left(X_0+K_S\,Y_{\mathrm{xs}}+S_0\,Y_{\mathrm{xs}}\right)}{X_m\,\left(X_0+S_0\,Y_{\mathrm{xs}}\right)}+\frac{K_S\,Y_{\mathrm{xs}}\,\ln(-S_0\,Y_{\mathrm{xs}})}{\left(X_0+S_0\,Y_{\mathrm{xs}}\right)\,\left(X_0-X_m+S_0\,Y_{\mathrm{xs}}\right)},X\right)$$

>> eqn7 = diff(S,t) == -mu_m*S*(1-(X0+Y_xs*(S0-S))/X_m)*(X0+Y_xs*(S0-S))/(K_S+ S)/Y_xs;

>> cond2 = S(0) == S0;

>> solS = dsolve(eqn7,cond2);

>> latex(eqn7)

ans =

$$\frac{\partial}{\partial t}S(t)=\frac{\mu_m\,S(t)\,\left(X_0+Y_{\mathrm{xs}}\,\left(S_0-S(t)\right)\right)\,\left(\frac{X_0+Y_{\mathrm{xs}}\,\left(S_0-S(t)\right)}{X_m}-1\right)}{Y_{\mathrm{xs}}\,\left(K_S+S(t)\right)}$$

solS =

(K_S*log(S/S0))/(S0^2*Y_xs^2 + 2*S0*X0*Y_xs - X_m*S0*Y_xs + X0^2 - X_m*X0) + (log((X0 - X_m)/(X0 - X_m - S*Y_xs + S0*Y_xs))*(X0 - X_m + Y_xs*(K_S + S0)))/(- X_m^2*Y_xs + S0*X_m*Y_xs^2 + X0*X_m*Y_xs) + (log((X0 - S*Y_xs + S0*Y_xs)/S0)*(X0 + Y_xs*(K_S + S0)))/(S0*X_m*Y_xs^2 + X0*X_m*Y_xs) - (mu_m*t)/(X_m*Y_xs)

>> latex(solS)

ans =

$$\frac{K_S\,\ln\left(\frac{S}{S_0}\right)}{{S_0}^2\,{Y_{\mathrm{xs}}}^2+2\,S_0\,X_0\,Y_{\mathrm{xs}}-X_m\,S_0\,Y_{\mathrm{xs}}+{X_0}^2-X_m\,X_0}+\frac{\ln\left(\frac{X_0-X_m}{X_0-X_m-S\,Y_{\mathrm{xs}}+S_0\,Y_{\mathrm{xs}}}\right)\,\left(X_0-X_m+Y_{\mathrm{xs}}\,\left(K_S+S_0\right)\right)}{-{X_m}^2\,Y_{\mathrm{xs}}+S_0\,X_m\,{Y_{\mathrm{xs}}}^2+X_0\,X_m\,Y_{\mathrm{xs}}}+\frac{\ln\left(\frac{X_0-S\,Y_{\mathrm{xs}}+S_0\,Y_{\mathrm{xs}}}{X_0}\right)\,\left(X_0+Y_{\mathrm{xs}}\,\left(K_S+S_0\right)\right)}{S_0\,X_m\,{Y_{\mathrm{xs}}}^2+X_0\,X_m\,Y_{\mathrm{xs}}}=\frac{\mu_m\,t}{X_m\,Y_{\mathrm{xs}}}$$

$$\frac{K_S\,\ln\left(\frac{S(t)}{S_0}\right)}{{S_0}^2\,{Y_{\mathrm{xs}}}^2+2\,S_0\,X_0\,Y_{\mathrm{xs}}-X_m\,S_0\,Y_{\mathrm{xs}}+{X_0}^2-X_m\,X_0}+\frac{\ln\left(\frac{X_0-X_m}{X_0-X_m+S_0\,Y_{\mathrm{xs}}-Y_{\mathrm{xs}}\,S(t)}\right)\,\left(X_0-X_m+Y_{\mathrm{xs}}\,\left(K_S+S_0\right)\right)}{-{X_m}^2\,Y_{\mathrm{xs}}+S_0\,X_m\,{Y_{\mathrm{xs}}}^2+X_0\,X_m\,Y_{\mathrm{xs}}}+\frac{\ln\left(\frac{X_0+S_0\,Y_{\mathrm{xs}}-Y_{\mathrm{xs}}\,S(t)}{S_0}\right)\,\left(X_0+Y_{\mathrm{xs}}\,\left(K_S+S_0\right)\right)}{S_0\,X_m\,{Y_{\mathrm{xs}}}^2+X_0\,X_m\,Y_{\mathrm{xs}}}-\frac{\mu_m\,t}{X_m\,Y_{\mathrm{xs}}}$$

$$\mathrm{solve}\left(\frac{K_S\,\ln(S)}{{S_0}^2\,{Y_{\mathrm{xs}}}^2+2\,S_0\,X_0\,Y_{\mathrm{xs}}-X_m\,S_0\,Y_{\mathrm{xs}}+{X_0}^2-X_m\,X_0}-\frac{\ln\left(X_0-X_m-S\,Y_{\mathrm{xs}}+S_0\,Y_{\mathrm{xs}}\right)\,\left(X_0-X_m+Y_{\mathrm{xs}}\,\left(K_S+S_0\right)\right)}{-{X_m}^2\,Y_{\mathrm{xs}}+S_0\,X_m\,{Y_{\mathrm{xs}}}^2+X_0\,X_m\,Y_{\mathrm{xs}}}+\frac{\ln\left(X_0-S\,Y_{\mathrm{xs}}+S_0\,Y_{\mathrm{xs}}\right)\,\left(X_0+Y_{\mathrm{xs}}\,\left(K_S+S_0\right)\right)}{S_0\,X_m\,{Y_{\mathrm{xs}}}^2+X_0\,X_m\,Y_{\mathrm{xs}}}=\frac{\ln\left(X_0\right)\,\left(X_0+Y_{\mathrm{xs}}\,\left(K_S+S_0\right)\right)}{S_0\,X_m\,{Y_{\mathrm{xs}}}^2+X_0\,X_m\,Y_{\mathrm{xs}}}+\frac{K_S\,\ln\left(S_0\right)}{{S_0}^2\,{Y_{\mathrm{xs}}}^2+2\,S_0\,X_0\,Y_{\mathrm{xs}}-X_m\,S_0\,Y_{\mathrm{xs}}+{X_0}^2-X_m\,X_0}-\frac{\ln\left(X_0-X_m\right)\,\left(X_0-X_m+Y_{\mathrm{xs}}\,\left(K_S+S_0\right)\right)}{-{X_m}^2\,Y_{\mathrm{xs}}+S_0\,X_m\,{Y_{\mathrm{xs}}}^2+X_0\,X_m\,Y_{\mathrm{xs}}}+\frac{\mu_m\,t}{X_m\,Y_{\mathrm{xs}}},S\right)$$

### 4. Matlab code to solve the hybrid Logistic-Monod cell growth model in CSTR

```
>> syms S X D mu_max X_max S_F K_S Y_xs mu
>> eqn1 = mu == mu_max*(1-X/X_max)*S/(K_S+S)
>> eqn2 = mu*X-D*X == 0
>> eqn3 = -mu*X/Y_xs +D*(S_F-S) == 0;
>> eqns =[eqn1,eqn2, eqn3];
```

>> vars = [mu S X];

>> [solmu, solS, solX] = solve(eqns, vars)

>> latex(eqn1)

ans =

$$\mu = -\frac{S\,\mu_{\mathrm{max}}\left(\frac{X}{X_{\mathrm{max}}} - 1\right)}{K_S + S}$$

>> latex(eqn2)

ans =

$$X\,\mu - D\,X = 0$$

>> latex(eqn3)

ans =

$$-D\,(S - S_F) - \frac{X\,\mu}{Y_{\mathrm{xs}}} = 0$$

>> latex(solS(2,1))

$$S_F - \frac{\sqrt{D^2\,X_{\mathrm{max}}^2 + 2\,D\,S_F\,X_{\mathrm{max}}\,Y_{\mathrm{xs}}\,\mu_{\mathrm{max}} - 2\,D\,X_{\mathrm{max}}^2\,\mu_{\mathrm{max}} + 4\,K_S\,D\,X_{\mathrm{max}}\,Y_{\mathrm{xs}}\,\mu_{\mathrm{max}} + S_F^2\,Y_{\mathrm{xs}}^2\,\mu_{\mathrm{max}}^2 - 2\,S_F\,X_{\mathrm{max}}\,Y_{\mathrm{xs}}\,\mu_{\mathrm{max}}^2 + X_{\mathrm{max}}^2\,\mu_{\mathrm{max}}^2} + X_{\mathrm{max}}\,\mu_{\mathrm{max}} - D\,X_{\mathrm{max}} + S_F\,Y_{\mathrm{xs}}\,\mu_{\mathrm{max}}}{2\,Y_{\mathrm{xs}}\,\mu_{\mathrm{max}}}$$

>> latex(solS(3,1))

$$S_F + \frac{\sqrt{D^2\,X_{\mathrm{max}}^2 + 2\,D\,S_F\,X_{\mathrm{max}}\,Y_{\mathrm{xs}}\,\mu_{\mathrm{max}} - 2\,D\,X_{\mathrm{max}}^2\,\mu_{\mathrm{max}} + 4\,K_S\,D\,X_{\mathrm{max}}\,Y_{\mathrm{xs}}\,\mu_{\mathrm{max}} + S_F^2\,Y_{\mathrm{xs}}^2\,\mu_{\mathrm{max}}^2 - 2\,S_F\,X_{\mathrm{max}}\,Y_{\mathrm{xs}}\,\mu_{\mathrm{max}}^2 + X_{\mathrm{max}}^2\,\mu_{\mathrm{max}}^2} - X_{\mathrm{max}}\,\mu_{\mathrm{max}} + D\,X_{\mathrm{max}} - S_F\,Y_{\mathrm{xs}}\,\mu_{\mathrm{max}}}{2\,Y_{\mathrm{xs}}\,\mu_{\mathrm{max}}}$$

$$S_F + \frac{\sqrt{D^2\,X_m^2 + 2\,D\,S_F\,X_m\,Y_{\mathrm{xs}}\,\mu_m - 2\,D\,X_m^2\,\mu_m + 4\,K_S\,D\,X_m\,Y_{\mathrm{xs}}\,\mu_m + S_F^2\,Y_{\mathrm{xs}}^2\,\mu_m^2 - 2\,S_F\,X_m\,Y_{\mathrm{xs}}\,\mu_m^2 + X_m^2\,\mu_m^2} - X_m\,\mu_m + D\,X_m - S_F\,Y_{\mathrm{xs}}\,\mu_m}{2\,Y_{\mathrm{xs}}\,\mu_m}$$

$$\frac{\sqrt{\left((D - \mu_m)X_m + S_F\,Y_{\mathrm{xs}}\,\mu_m\right)^2 + 4\,K_S\,D\,X_m\,Y_{\mathrm{xs}}\,\mu_m} - X_m\,\mu_m + D\,X_m + S_F\,Y_{\mathrm{xs}}\,\mu_m}{2\,Y_{\mathrm{xs}}\,\mu_m}$$

$$\frac{\sqrt{(X_m\,(D - \mu_m) + S_F\,Y_{\mathrm{xs}}\,\mu_m)^2 + 4\,D\,K_S\,X_m\,Y_{\mathrm{xs}}\,\mu_m} - X_m\,\mu_m + D\,X_m + S_F\,Y_{\mathrm{xs}}\,\mu_m}{2\,Y_{\mathrm{xs}}\,\mu_m}$$

>> latex(solX(2,1))

$$\frac{\sqrt{D^2\,X_{\mathrm{max}}^2 + 2\,D\,S_F\,X_{\mathrm{max}}\,Y_{\mathrm{xs}}\,\mu_{\mathrm{max}} - 2\,D\,X_{\mathrm{max}}^2\,\mu_{\mathrm{max}} + 4\,K_S\,D\,X_{\mathrm{max}}\,Y_{\mathrm{xs}}\,\mu_{\mathrm{max}} + S_F^2\,Y_{\mathrm{xs}}^2\,\mu_{\mathrm{max}}^2 - 2\,S_F\,X_{\mathrm{max}}\,Y_{\mathrm{xs}}\,\mu_{\mathrm{max}}^2 + X_{\mathrm{max}}^2\,\mu_{\mathrm{max}}^2} + X_{\mathrm{max}}\,\mu_{\mathrm{max}} - D\,X_{\mathrm{max}} + S_F\,Y_{\mathrm{xs}}\,\mu_{\mathrm{max}}}{2\,\mu_{\mathrm{max}}}$$

>> latex(solX(3,1))

$$-\frac{\sqrt{D^2\,X_{\mathrm{max}}^2 + 2\,D\,S_F\,X_{\mathrm{max}}\,Y_{\mathrm{xs}}\,\mu_{\mathrm{max}} - 2\,D\,X_{\mathrm{max}}^2\,\mu_{\mathrm{max}} + 4\,K_S\,D\,X_{\mathrm{max}}\,Y_{\mathrm{xs}}\,\mu_{\mathrm{max}} + S_F^2\,Y_{\mathrm{xs}}^2\,\mu_{\mathrm{max}}^2 - 2\,S_F\,X_{\mathrm{max}}\,Y_{\mathrm{xs}}\,\mu_{\mathrm{max}}^2 + X_{\mathrm{max}}^2\,\mu_{\mathrm{max}}^2} - X_{\mathrm{max}}\,\mu_{\mathrm{max}} + D\,X_{\mathrm{max}} - S_F\,Y_{\mathrm{xs}}\,\mu_{\mathrm{max}}}{2\,\mu_{\mathrm{max}}}$$

$$-\frac{\sqrt{\left((D-\mu_m)X_m+S_F Y_{xs}\mu_m\right)^2+4K_S D X_m Y_{xs}\mu_m}-X_m\mu_m+D X_m-S_F Y_{xs}\mu_m}{2\mu_m}$$

$$\frac{X_m\mu_m-\sqrt{(X_m(D-\mu_m)+S_F Y_{xs}\mu_m)^2+4 D K_S X_m Y_{xs}\mu_m}-D X_m+S_F Y_{xs}\mu_m}{2\mu_m}$$

### 5. Matlab code to generate Figure 2

```
syms K_S D S Y_xs mu_m X_m S_F X
S1 = K_S*D/(mu_m-D);
X1 = Y_xs*(S_F-K_S*D/(mu_m-D));
subplot(1,3,1)
fplot(subs(S1, [mu_m, X_m, S_F, K_S, Y_xs], [1.6, 10, 20, 1, 0.8]), [0 1.6] , 'LineWidth',2.0)
hold on
fplot(subs(X1, [mu_m, X_m, S_F, K_S, Y_xs], [1.6, 10, 20, 1, 0.8]), [0 1.6] , 'LineWidth',2.0)
fplot(subs(S1, [mu_m, X_m, S_F, K_S, Y_xs], [1.6, 10, 20, 1, 0.3]), [0 1.6] , 'LineWidth',2.0)
fplot(subs(X1, [mu_m, X_m, S_F, K_S, Y_xs], [1.6, 10, 20, 1, 0.3]), [0 1.6] , 'LineWidth',2.0)
>> xlabel('Dilution rate (1/h)')
ylabel('Cell and substrate (g/L)')
legend('Y_{xs}=0.8', 'Y_{xs}=0.8', 'Y_{xs}=0.3', 'Y_{xs}=0.3')
title('(a)');
ylim([0 16])
legend boxoff
ax = gca;
ax.FontSize = 16



>> syms S2 X2
>> S2 = (D*X_m-X_m*mu_m+S_F*Y_xs*mu_m)/Y_xs/mu_m;
>> X2 = X_m*(1-D/mu_m);
>> subplot(1,3,2)
fplot(subs(S2, [mu_m, X_m, S_F, K_S, Y_xs], [1.6, 10, 20, 1, 0.8]), [0 1.6] , 'LineWidth',2.0)
hold on
```

```
fplot(subs(X2, [mu_m, X_m, S_F, K_S, Y_xs], [1.6, 10, 20, 1, 0.8]), [0 1.6] , 'LineWidth',2.0)

fplot(subs(S2, [mu_m, X_m, S_F, K_S, Y_xs], [1.6, 10, 20, 1, 0.3]), [0 1.6] , 'LineWidth',2.0)

fplot(subs(X2, [mu_m, X_m, S_F, K_S, Y_xs], [1.6, 10, 20, 1, 0.3]), [0 1.6] , 'LineWidth',2.0)

>> xlabel('Dilution rate (1/h)')

ylabel('Cell and substrate (g/L)')

legend('Y_{xs}=0.8', 'Y_{xs}=0.8', 'Y_{xs}=0.3', 'Y_{xs}=0.3')

title('(b)');

ylim([0 16])

legend boxoff

ax = gca;

ax.FontSize = 16


>> syms S3 X3

>> S3 = (sqrt(((D-mu_m)*X_m+S_F*Y_xs*mu_m)^2+4*K_S*D*X_m*Y_xs*mu_m)-X_m*mu_m+D*X_m+S_F*Y_xs*mu_m)/(2*Y_xs*mu_m);

>> X3 = -(sqrt(((D-mu_m)*X_m+S_F*Y_xs*mu_m)^2+4*K_S*D*X_m*Y_xs*mu_m)-X_m*mu_m+D*X_m-S_F*Y_xs*mu_m)/(2*mu_m);

>> subplot(1,3,3)

fplot(subs(S3, [mu_m, X_m, S_F, K_S, Y_xs], [1.6, 10, 20, 1, 0.8]), [0 1.6] , 'LineWidth',2.0)

hold on

fplot(subs(X3, [mu_m, X_m, S_F, K_S, Y_xs], [1.6, 10, 20, 1, 0.8]), [0 1.6] , 'LineWidth',2.0)

fplot(subs(S3, [mu_m, X_m, S_F, K_S, Y_xs], [1.6, 10, 20, 1, 0.3]), [0 1.6] , 'LineWidth',2.0)

fplot(subs(X3, [mu_m, X_m, S_F, K_S, Y_xs], [1.6, 10, 20, 1, 0.3]), [0 1.6] , 'LineWidth',2.0)

>> xlabel('Dilution rate (1/h)')

ylabel('Cell and substrate (g/L)')

legend('Y_{xs}=0.8', 'Y_{xs}=0.8', 'Y_{xs}=0.3', 'Y_{xs}=0.3')

title('(c)');

ylim([0 16])

legend boxoff

ax = gca;

ax.FontSize = 16
```
